# *Clostridioides difficile* in Equidae necropsied in northwestern France, between 2019 and 2021

**DOI:** 10.1101/2024.09.30.615820

**Authors:** Sandrine Petry, Jackie Tapprest, Karine Maillard, Frédéric Barbut, Fabien Duquesne, Sofia Kozak, Nathalie Foucher, Maud Bernez-Romand, Ludovic Bridoux, Isabelle Poquet

## Abstract

*Clostridioides difficile* is an anaerobic, spore-forming enteropathogen, which is much more extensively studied in humans than animals, despite growing evidence supporting its importance in One Health. We evaluated *C. difficile* occurrence, diversity, circulation and virulence in French Equidae (n=100) after their necropsy in northwestern France (Normandy), from 2019 to 2021. We systematically recovered all cecal contents and any watery (possibly diarrheal) intestinal contents. We isolated *C. difficile* strains, determined their toxin gene profile by PCR and established their PCR-ribotype according to the WEBRIBO database. We also performed free toxin detection.

Twenty-seven Equidae were positive for *C. difficile* and twenty had a toxigenic strain, including one animal also colonized by a non-toxigenic strain. Toxigenic isolates belonged to eight common ribotypes: i) 078 and 126 (*tcdA^+^ tcdB^+^ cdt^+^* according to multiplex PCR, i.e. able to produce toxin A, toxin B and binary toxin), ii) 005, 012, 020, AI-53 and FR227 (*tcdA^+^ tcdB^+^*), and iii) 017 (*tcdB^+^*). Non-toxigenic isolates were of ribotypes 009, 035 and 439. Ribotypes 017 and 009 were predominant (n=5). In two cases, Equidae of the same premises shared the same ribotype, either 020 or 009. Free toxins were detected in four animals: they displayed signs of diarrhea and a *C. difficile* of ribotype 126 (n=1) or 017 (n=3) as the only detected enteropathogen, suggesting a *C. difficile* infection (CDI). Three of them had received antibiotics. Two had died from an entero-toxic infection, for which *C. difficile* ribotype 017 was the only identified cause.

French Equidae were found to display common pathogenic *C. difficile*, and ribotype 017 was highly virulent. These findings are of concern from a One Health perspective.

**Importance:** *C. difficile*, a major enteropathogen widely disseminated in the environment, is a One Health issue. Animals are raising concern as human contamination sources. Equidae are in close contact with humans and also develop *post*-antibiotic and healthcare-associated CDIs. The systematic survey of Equidae necropsied from 2019 to 2021 in the leading region for horse breeding in France, revealed that 20% harbored pathogenic strains. These belonged to clinically important ribotypes, raising the possibility of cross-species, possibly zoonotic transmission. Free toxins, which are rarely tested in animals, were detected in four animals with signs of diarrhea and a toxigenic *C. difficile* as the only identified enteropathogen, suggesting CDI. In two of them, *C. difficile* ribotype 017 was the only identified cause of entero-toxic disease and death.

French Equidae could play a role in the dissemination of pathogenic *C. difficile* and notably ribotype 017. They should be surveilled carefully from a One Health perspective.

## Introduction

*Clostridioides difficile* is a spore-forming anaerobe responsible for diarrhea and colitis (1) *C. difficile,* whose spores are transmitted by the fecal-oral route, is a major cause of healthcare-associated diarrhea and a rising cause of community-acquired infections (1). *C. difficile* infections (CDIs) can be severe, with complications and recurrences, and life threatening. Toxin A (TcdA) and Toxin B (TcdB) are responsible for symptoms, with TcdB alone being sufficient to cause disease. The binary toxin (CDT) has an accessory role (2). Non-toxigenic strains are unable to cause disease, while toxigenic strains can cause disease or be asymptomatically carried (3). Antibiotics, hospitalization and age increase the risk of developing CDI (1). Antibiotics notably induce microbiota dysbiosis, which favors spore germination, vegetative multiplication, and ultimately, toxin production (1). Healthcare-associated CDIs are treated with fidaxomicin and vancomycin (4), which can contribute to microbiota dysbiosis and favor *C. difficile* re-emergence, possibly from persisting biofilms (5, 6). In this context, the last resort treatment is fecal microbiota transplant (FMT) (4).

*C. difficile* is a One Health issue (7–10). Spores, which are highly resistant, are widely disseminated by fecal spreading in the environment and can contaminate different hosts (8, 9, 11–13). *C. difficile* strains can be transmitted between host species (10, 14–16). Toxigenic strains can cause diarrhea (CDI) in a wide range of wild, farm and companion animals (8, 9, 11–13). Equidae, which are monogastric mammals, are closely interacting with humans and relevant to zoonotic and anthropo-zoonotic transmission. Importantly, in these animals of high economic value (https://www.ifce.fr), antibiotics treatments and hospitalization increase the risk of developing CDI, as it is the case in humans. In Equidae, *C. difficile* is also responsible for outbreaks and sporadic cases of CDI (17–21). Both Equidae and humans are treated with vancomycin and have long been treated with metronidazole, which is still used in animals but replaced with fidaxomicin for hospitalized patients. Like humans, Equidae can also be treated with FMT (22). The main difference between Equidae and humans as *C. difficile-*infected populations is age. While <3-years-old children are mainly asymptomatic carriers (3), CDI is common in foals (17, 18).

Data about the importance of *C. difficile* in horses (7, 9, 11, 19, 23) remain scarce in France. *C. difficile* testing is not systematic, and the detection of a toxigenic strain is not sufficient to diagnose CDI (3). Here, we studied *C. difficile* in Equidae necropsied in Normandy, the leading region for foal birth in France (12,600 foals and 145,000 Equidae in 2021, according to the French Institute for Horse and Riding (IFCE) https://www.ifce.fr and https://statscartes.ifce.fr/storage/files/1/pdf/IFCE-Depliant-chiffres-cles-2022-WEB.pdf). Necropsy (24) provided the opportunity both to recover digestive contents from a population with diverse clinical pictures, and to get access to detailed animal data including the cause of animal death. Our objective was to *post-mortem* evaluate *C. difficile* occurrence, diversity, putative circulation and virulence in French Equidae.

## Materials and methods

### Equine necropsy

At the French National Agency for Food, Environmental and Occupational Health & Safety (ANSES), the PhEED Unit in Normandy is in charge of surveillance, prevention and study of equine diseases. It notably performs equine necropsies at the request of breeders or veterinarians to determine the cause of animal death. The standardized procedure includes examination of the cadaver, evisceration and organ examination. Animal identification data, previous treatments, including antibiotics in the two-three weeks before death, necropsy observations, pathogen identification and the cause of animal death, established by the veterinarian in charge of the necropsy, are registered (https://sitesv2.anses.fr/fr/minisite/resumeq/presentation-de-resumeq) (24).

In 2018, we chose five necropsied Equidae with *post-mortem* signs of endo-enterotoxaemia and diarrhea and collected their watery intestinal contents for a pilot study. Then, from May 2019 to August 2021, we studied all one hundred necropsied animals (except fetus and stillborn foals), irrespective of clinical picture or cause of death. This total number, after two years characterized by Covid-19 and lockdowns, was relatively low, matching the previous annual average. Animal digestive contents were systematically recovered in the caecum (CAE; n=100), and in case of a watery consistency, in any other intestine segment (CDL; n=30), notably the colon. Aliquots of all digestive contents except a cecal one (129 samples in 2019-2021) were preserved at 4°C, −20°C and −70°C.

This study followed institutional guidelines and was approved by the scientific committee of IFCE, the French Institute on Horse and Riding (https://www.ifce.fr) that funded the project. All data are anonymous.

### Standard pathogen testing

In case of *post-mortem* observed signs of infection, the veterinarian requested routine pathogen testing in relevant samples (Table S1). For intestinal infections, usual testing in digestive contents was as follows. *Escherichia coli* colonies, after detection on Eosin Methylene Blue plates, were streaked on blood plates to see hemolysis. *Salmonella* spp. were isolated on XLD plates (Thermo Scientific). *C. difficile* was detected on ChromID (BioMérieux) under anaerobiosis, and *Clostridium perfringens* and *Paeniclostridium sordellii* on Columbia agar containing 5% sheep blood (BioMérieux). Matrix-Assisted Laser Desorption/Ionization Time-Of-Flight (MALDI-TOF) Mass Spectrometry (MS) (MBT Smart mass spectrometer, Compass and FlexControl softwares, Bruker Daltonics) confirmed identification. Rotavirus was identified by PCR. Intestinal parasite communities were identified and quantified by visual inspection (25).

### *C. difficile* testing, isolation and characterization

*C. difficile* presence was specifically and independently tested in all preserved contents, irrespective of whether the veterinarian observed signs of intestinal infections and requested enteropathogen testing or not. A sample of any digestive content was streaked onto ChromID or CLO plates (BioMérieux) and incubated for 48h at 37°C under anaerobiosis. In case of a negative result, a 1g-aliquot was harvested after sample homogenization, inoculated in rich, permissive medium: BHI broth supplemented by cysteine (0.1%), yeast extract (5 g/L), taurocholate (1 g/L), D-cycloserine (250 µg/mL) and cefoxitin (8 µg/mL) (OXOID), and finally incubated for 3-7 days (enrichment) before plating onto ChromID. A positive stool was included as a control. Suspicious, irregular and/or black colonies were screened by PCRs targeting *tpi*, *tcdA* and *tcdB* genes (26, 27). Identification was confirmed by MALDI-TOF MS. For each confirmed isolate, a single colony per positive sample was grown in BHI broth at 37°C under anaerobiosis to be preserved in glycerol (17%) at −70°C, except in two cases: two strains (non-toxigenic in animal #48 and toxigenic in #54) characterized without being preserved, could not be recovered later.

All preserved isolates (N=34 from n=25 Equidae in 2019-2021 and N=1 in 2018) were characterized by multiplex PCR targeting: i) *tpi* gene, ii) wild-type or truncated *tcd* and *cdt* genesand iii) *lok* fragment characterizing non-toxigenic strains, followed by capillary gel electrophoresis (28). For ribotyping, PCR profiles after capillary gel-based electrophoresis (29) were analyzed with GeneMapper software (Thermo Fischer Scientific, Villebon-sur-Yvette, France) and ribotypes assigned according to database WEBRIBO (https://webribo.ages.at/). The first isolates were characterized by Multi Locus Sequence Typing (MLST) (30).

### Susceptibility to Vancomycin and to metronidazole

The Minimum Inhibitory Concentration (MIC) was determined using Etest (BioMérieux) according to the EUCAST and CA-SFM recommendations in 2021 (https://www.sfm-microbiologie.org/wp-content/uploads/2021/04/CASFM2021V1.0.AVRIL_2021.pdf). Briefly, a suspension of 1.0 MacFarland was prepared in Schaedler broth and spread on a pre-reduced Brucella blood agar (Becton Dickinson) supplemented with vitamin K1 (10□mg/L), hemin (5□mg/L) and defibrinated horse blood (5%) (OXOID). Of note, after we achieved our experiments, Brucella blood agar medium had been published to be suboptimal. A strip, in which antibiotic concentration ranged from 0.0016 to 256mg/L, was laid on the plates, which were then incubated for 24□h (vancomycin) or 48h (metronidazole)□at 37°C under anaerobiosis (using Thermo Scientific™ Oxoid™ AnaeroGen™ Compact sachets). We used *C. difficile* reference strain ATCC70075 as a quality control.

### Toxin detection

1. *C. difficile* toxin A, toxin B and glutamate dehydrogenase (GDH) were detected in digestive contents using an Enzymatic Immuno-Assay (Quick Check Complete, Alere), according to manufacturer’s instructions. Every time a new kit was opened, the supplied positive control was tested. For each series of experiments, we included, as a positive control, an aliquot of a positive human stool stored at −80°C. A second independent observation was performed to confirm any weak positive result (band of weaker intensity than that of the control).

### Statistical analysis

We used a Fisher’s exact test to analyze the equine population composition, after excluding unknown values, and a Mann-Whitney test to compared strain MICs.

## Results

### Necropsied Equidae

We studied all Equidae (except fetuses and stillborn foals) (n=100) necropsied at ANSES in Normandy from May 2019 to August 2021, without any selection (Table S1). All animals except three (n=97) originated from Normandy, two from Brittany and Pays de Loire and one from an unknown region. Most animals were females (67%) and <1-year-old animals (59%), with foals (<6-months) representing half of the population (48%) (Table 1). Nine breeds were present, mostly Thoroughbred (42%), Trotters (31%) and Saddlebred horses (11%) (Table 1). Animals had died from very diverse (>30) causes including infectious or non-infectious diseases and trauma (Table S1). The first and second most prevalent causes of death were i) endo-enterotoxaemia (12%), an enteric disease caused by toxin-producing bacteria like Salmonella and Clostridia (20), and ii) infection by *Rhodococcus equi* (9%), common in foals (31).

**Table 1.** Equine population: data summary. This contingency table shows the composition of the equine population (number of necropsied animals). In lines, Equidae are categorized according to: i) necropsy year, ii) identity data (sex, age, breed), iii) *post-mortem* observations (signs of diarrhea or of endo-enterotoxaemia), iv) *ante-mortem* data (previous antibiotic treatment or hospitalization) (Table S1). In columns, Equidae are categorized according to *C. difficile*: i) its presence or not, assessed in the whole equine population (n=100), ii) *C. difficile* pathogenicity or not, in the subpopulation of *C. difficile*-positive animals (n=27) and finally iii) its production of toxins (virulence) or not, in the subpopulation of animals with a toxigenic *C. difficile* (n=20). *the information was missing in the necropsy register (Table S1). In these cases, the number of animals is indicated in italics for full and complete information, but these animals were excluded from the statistical analysis using Fisher’s exact test. Statistical significance: P-values < 0.05, which are shown in bold, on a grey background.

| Equidae | *C. difficile* |  |  | Pathogenicity |  |  |  | Toxin production |  |  |
| --- | --- | --- | --- | --- | --- | --- | --- | --- | --- | --- |
|  | No | Yes | P-value | Non-toxicogenic | Toxigenic | Both | P-value | No | Yes | P-value |
| **Necropsy Year** |  |  |  |  |  |  |  |  |  |  |
| 2019 | 32 | 15 | 0.60 | 3 | 11 | 1 | 0.91 | 8 | 4 | 0.30 |
| 2020 | 28 | 8 |  | 3 | 5 | 0 |  | 5 | 0 |  |
| 2021 | 13 | 4 |  | 1 | 3 | 0 |  | 3 | 0 |  |
| **Animal data** |  |  |  |  |  |  |  |  |  |  |
| **Sex** |  |  |  |  |  |  |  |  |  |  |
| Female | 46 | 21 | 0.23 | 7 | 13 | 1 | 0.34 | 11 | 3 | 1.00 |
| Male, gelding | 26 | 6 |  | 0 | 6 | 0 |  | 5 | 1 |  |
| Unknown* | 1 | 0 |  |  |  |  |  |  |  |  |
| **Age** |  |  |  |  |  |  |  |  |  |  |
| Foal | 33 | 15 | 0.55 | 6 | 9 | 0 | 0.42 | 6 | 3 | 0.79 |
| Yearling | 10 | 1 |  | 0 | 1 | 0 |  | 1 | 0 |  |
| Young | 5 | 2 |  | 0 | 2 | 0 |  | 2 | 0 |  |
| Adult | 25 | 9 |  | 1 | 7 | 1 |  | 7 | 1 |  |
| **Breed** |  |  |  |  |  |  |  |  |  |  |
| Thoroughbred | 30 | 12 | 0.80 | 6 | 6 | 0 | 0.15 | 6 | 0 | **0.038** |
| Trotter | 23 | 8 |  | 1 | 6 | 1 |  | 3 | 4 |  |
| Saddlebred horse | 9 | 2 |  | 0 | 2 | 0 |  | 2 | 0 |  |
| Other breeds | 9 | 5 |  | 0 | 5 | 0 |  | 5 | 0 |  |
| Unknown* | 2 | 0 |  |  |  |  |  |  |  |  |
| Thoroughbred / all other breeds | 30 / 41 | 12 / 15 | 1.0 | 6 / 1 | 6 / 13 | 0 / 1 | **0.026** | 6 / 10 | 0 / 4 | 0.27 |
| Trotter / all other breeds | 23 / 48 | 8 / 19 | 1.0 | 1 / 6 | 6 / 13 | 1 / 0 | 0.29 | 3 / 13 | 4 / 0 | **0.007** |
| **Post-mortem observed signs of** |  |  |  |  |  |  |  |  |  |  |
| **Diarrhea** |  |  |  |  |  |  |  |  |  |  |
| One sample per animal: CAE | 59 | 11 | **0.0002** | 3 | 8 | 0 | 1.0 | 8 | 0 | 0.12 |
| Two samples per animal: CAE + CDL | 14 | 16 |  | 4 | 11 | 1 |  | 8 | 4 |  |
| **Endo-enterotoxemia** |  |  |  |  |  |  |  |  |  |  |
| Yes | 7 | 13 | **0.00008** | 3 | 9 | 1 | 1.0 | 7 | 3 | 0.58 |
| No | 65 | 14 |  | 4 | 10 | 0 |  | 9 | 1 |  |
| Unknown* | 1 | 0 |  |  |  |  |  |  |  |  |
| **Risk factors** |  |  |  |  |  |  |  |  |  |  |
| **Previous antibiotic treatment** |  |  |  |  |  |  |  |  |  |  |
| Yes | 24 | 11 | 0.48 | 5 | 6 | 0 | 0.18 | 3 | 3 | **0.025** |
| No | 43 | 14 |  | 2 | 11 | 1 |  | 12 | 0 |  |
| Unknown* | 6 | 2 |  | 0 | 2 | 0 |  | 1 | 1 |  |
| **Previous hospitalization** |  |  |  |  |  |  |  |  |  |  |
| Yes | 22 | 11 | 0.47 | 4 | 7 | 0 | 0.65 | 7 | 0 | 0.25 |
| No | 49 | 16 |  | 3 | 12 | 1 |  | 9 | 4 |  |
| Unknown* | 2 | 0 |  | 3 | 12 | 1 |  | 9 | 4 |  |

### *C. difficile* prevalence

All cecal contents and any watery intestinal contents, possibly indicating diarrhea, were recovered. All except one were preserved (99 CAE and 30 CDL) and tested for *C. difficile*, directly or after enrichment. Candidate colonies were screened by standard PCRs (26, 27) and confirmed by MALDI-TOF. Twenty-seven animals were positive for *C. difficile*, in their two samples in nine cases (Table S1). *C. difficile* presence in Equidae was independent of animal sex, age, breed, antibiotic treatment or hospitalization, but significantly correlated to signs of diarrhea or endo-enterotoxaemia (Table 1).

### Pathogenicity

Thirty-four *<u>C</u>. <u>d</u>ifficile* isolates were preserved from twenty-five <u>E</u>quidae and named ‘*Cd*E’ (Table 2). They form the first French library of *C. difficile* strains originating from Equidae, CloDifEqui (Table 2). Their pathogenicity profile was characterized by multiplex PCR, which revealed non-toxigenic isolates (N=9) and toxigenic ones (N=25) with three profiles (*tcdA^+^ tcdB^+^ cdt^+^*, *tcdA^+^ tcdB^+^* and *tcdB^+^*) (Table 2). The former isolates originated from seven animals, including six Thoroughbred, and the latter from nineteen animals of six breeds (Table S1). One animal (#29) was co-colonized by a non-toxigenic strain (CAE) and a toxigenic one (CDL). In the eight other animals with two isolates, both were either non-toxigenic (#18, #60) or toxigenic with the same profile (#3, #32, #36, #37, #42, #43) (Table 2).

**Table 2.**
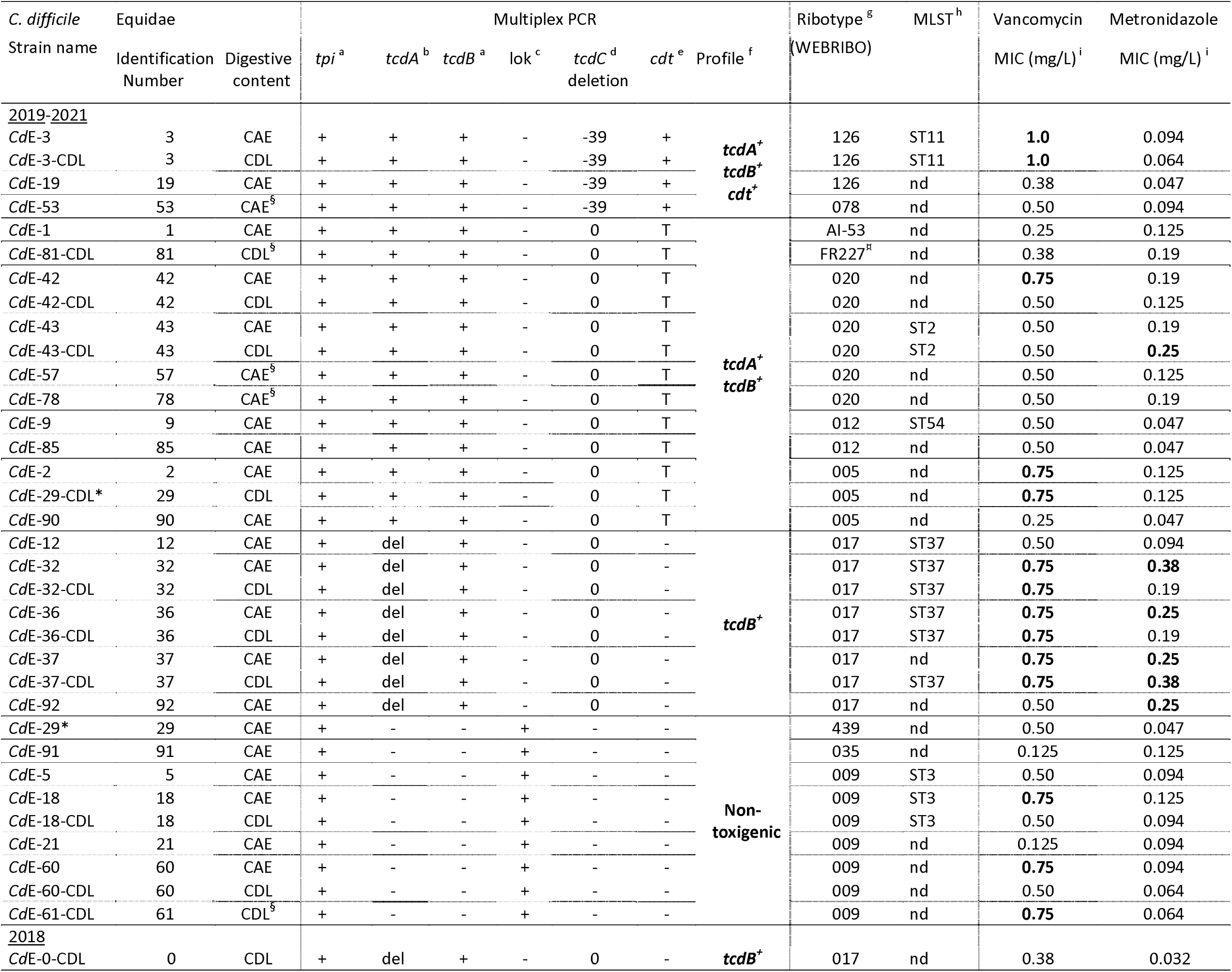
*C. difficile* strains isolated from necropsied Equidae. All preserved strains are listed. Their origin (animal and digestive content) is indicated and whether they have been isolated after enrichment is specified. For each strain, all results of molecular characterization: multiplex PCR results, PCR-ribotype and Multi Locus Sequence Type (MLST, Table S3), and the Minimal Inhibitory Concentration (MIC) of vancomycin and metronidazole are shown. ^a^PCR-fragment of wild-type size (‘+’) or absence of amplification (‘-’) ^b^PCR-fragments of wild-type size (‘+’) or of shorter size indicative of a deletion (‘del’) or absence of amplification (‘-’) ^c^PCR-fragment of 117 bp (‘+’, i. e. Pathogenicity Locus deletion and non-toxigenic strain) or no amplification (‘-’) ^d^39bp-deletion (‘-39’) or absence of deletion (‘0’, i. e. PCR-fragment of wild-type size) or absence of amplification (‘-’) ^e^PCR-fragment of wild-type size (‘+’) or truncated PCR-fragment (‘T’, i. e. presence of a pseudogene) or absence of amplification (‘-’) ^f^toxin gene profile according to multiplex PCR results: toxin genes of wild-type size or non-toxigenic ^g^PCR-ribotype assigned according to the database WEBRIBO ^h^MLST: Multi-Locus Sequence Type; nd: not determined ^i^MIC: Minimum Inhibitory concentration; the two highest values are in bold *the two strains co-colonizing the same animal ^§^the single digestive content (among the two ones recovered from the animal) from which *C. difficile* was isolated ^¤^Ribotyping was performed twice, as the initial automatic assignment according to database WEBRIBO was inconsistent with *cdtA* absence, according to multiplex PCR (a confirmed result): capillary gel-based electrophoresis revealed a double peak around 425bp, leading to the final assignation to ribotype FR227.

### Ribotype diversity

CloDifEqui isolates were assigned to eleven ribotypes and four clades (1, 2, 4 and 5) (Table 2, Table S2) (29). Toxigenic isolates were of eight ribotypes: 126 and 078 (*tcdA^+^ tcdB^+^ cdt^+^* and same *tcdC* truncation), 020, 005, 012, AI-53 and FR227 (*tcdA^+^ tcdB^+^*), and 017 (*tcdB^+^*). Non-toxigenic isolates belonged to ribotypes 009, 035 and 439. The two most prevalent ribotypes (n=5 animals) were 017 (N=8 isolates) and 009 (N=7).

The two strains of animal #29, which were toxigenic and non-toxigenic, respectively belonged to ribotype 005 and 439. In all other pairs of isolates originating from the same animal, both were of the same ribotype (Table 2).

### Susceptibility to metronidazole and to vancomycin

All strains were susceptible to the antibiotics used to treat CDI in Equidae, metronidazole and vancomycin (MICs lower than EUCAST breakpoints: 2 mg/L in both cases; https://www.eucast.org/fileadmin/src/media/PDFs/EUCAST_files/Breakpoint_tables/v_15.0_Breakpoint_Tables.pdf) (Table 2). For metronidazole, strain MICs ranged from 0.032 to 0.38mg/L, with a median value of 0.13 mg/L. For vancomycin, strain MICs ranged from 0.13 to 1 mg/L, with a median value of 0.5 mg/L. Strains of ribotype 017 and 126 showed the highest MIC for metronidazole (0.38 mg/L) and for vancomycin (1.0 mg/L), respectively.

### Co-localization of Equidae sharing the same ribotype

We cross-analyzed Equidae localization and *C. difficile* ribotypes (Figure 1, Table S1). In two premises between Caen and Pont-Audemer, some animals shared the same ribotype: i) 020, for two co-localized Equidae necropsied in 2019 (#42 and #43), and ii) 009 for four co-localized Equidae necropsied in 2019 (#5, #18 and #21) and 2020 (#61). This suggested that in each premises, the isolates of the same ribotype might be clonal (belong to a clonal complex) and that transmission might have occurred between animals or from a common environmental source.

**Figure 1.**
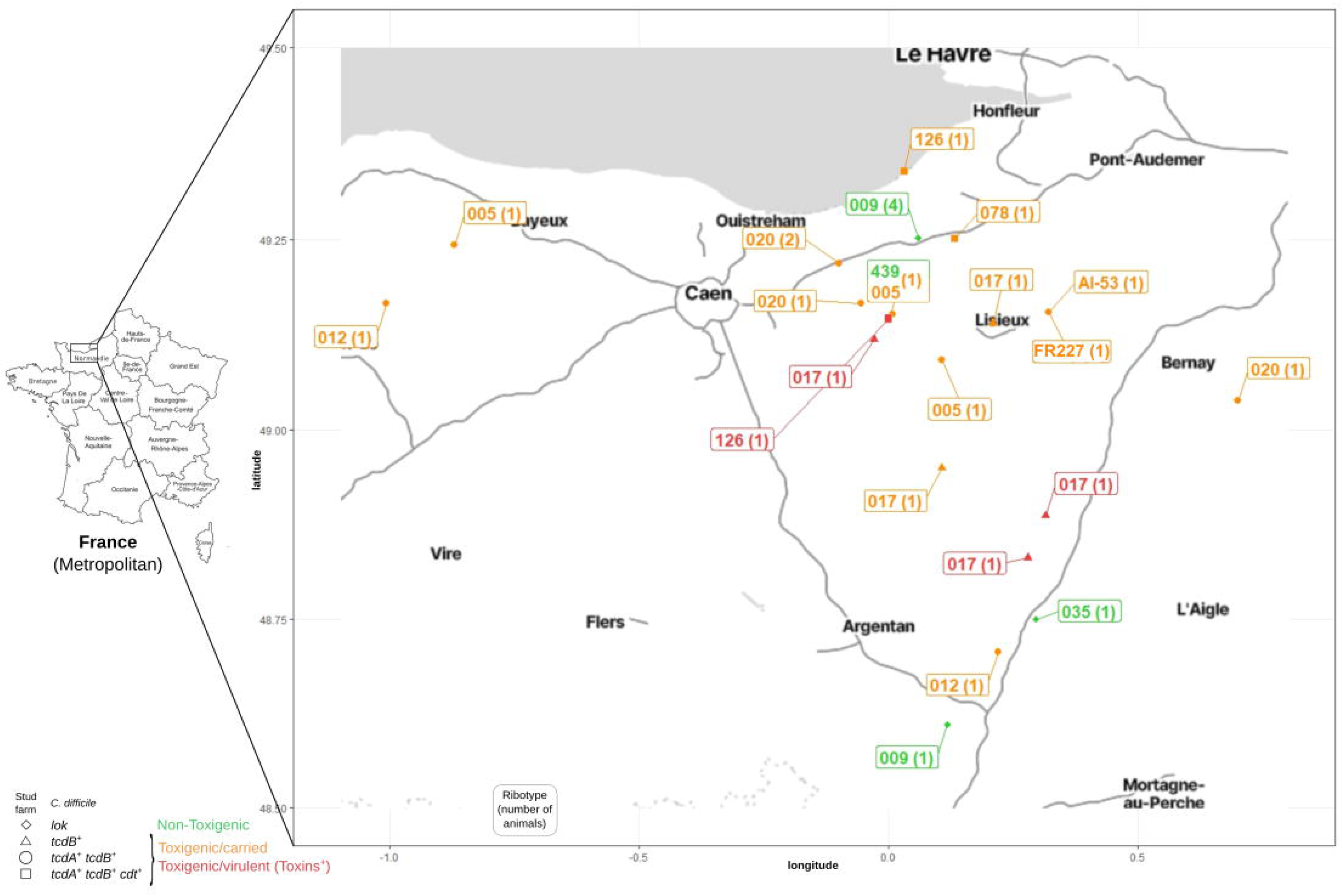
Geographical localization of *C. difficile*-positive Equidae. All *C. difficile*-positive Equidae are shown on a map of Normandy, according to the latitude and longitude coordinates of their stud farm (Table S1). Each animal/stud farm is indicated by a symbol depending on *C. difficile* and its toxin genes (multiplex PCR profile): squares for *tcdA^+^ tcdB^+^ cdt^+^*, circles for *tcdA^+^ tcdB^+^*, triangles for *tcdB^+^* and diamonds for non-toxigenic strains (Table 2). *C. difficile* ribotype (Table 2) is indicated in a box, and when several animals harbored this ribotype in the same premises (Table S1), their number appears between brackets. *C. difficile* pathogenicity and virulence are represented using a color code: green for non-toxigenic, orange for toxigenic and carried, and finally red for toxigenic and virulent (having produced toxins)..

### Toxin production

We tested the presence of free toxins in equine digestive contents using a standard enzymatic immuno-assay, developed to detect free toxin in human fecal material. To check the assay specificity when applied to these contents rather than human stools, we included samples expected to be toxin-negative: i) all ten samples displaying a non-toxigenic *C. difficile* (Table 3) and ii) twenty-seven *C. difficile*-negative samples (fifteen CDL and all samples from animals with endo-enterotoxaemia signs) (Table 3, Table S1, Table S3). No toxins could be detected in any of them, while GDH was positive in seven of the former (Table 3, Table S3).

**Table 3.** Free toxin detection. The results of free toxin detection are shown for the digestive contents of all *C. difficile*-positive Equidae. For each Equidae, identification number and the nature of recovered digestive content(s) (CAE and possibly CDL, in bold) are indicated. For each digestive content, *C. difficile* strain and pathogenicity are shown together with the results of toxin and GDH detection (enzymatic immuno-assay). Animals are listed according to *C. difficile* pathogenicity and to their diarrhea signs (watery CDL content). ^a^Strain name or ‘not preserved’ (an initially detected strain that could not be recovered later) or ‘none’ (absence of *C. difficile* detection) ^b^Method used to isolate the strain: Enrichment (‘+’) or not (‘-’, i. e. direct inoculation on plates) ^c^Toxigenic (in bold) or non-toxigenic strain ^d^Detection (‘+’) or absence of detection (‘-’); ‘weak’ indicates the intensity of detection was low compared to that obtained for the positive control (such a result was confirmed by two independent observations) NA: not applicable ND: not determined *co-colonized animal

| Equidae number | Digestive content | Strain <sup>a</sup> | Enrichment <sup>b</sup> | Pathogenicity <sup>c</sup> | Ribotype | GDH <sup>d</sup> | Free toxins <sup>d</sup><br>Toxins A and B |
| --- | --- | --- | --- | --- | --- | --- | --- |
| **3** | CAE | CdE-3 | - | **Toxigenic** | 126 | + | + |
|  | **CDL** | CdE-3-CDL | - | **Toxigenic** | 126 | + | + |
| **32** | CAE | CdE-32 | - | **Toxigenic** | 017 | + | + (weak) |
|  | **CDL** | CdE-32-CDL | - | **Toxigenic** | 017 | + | + (weak) |
| **36** | CAE | CdE-36 | - | **Toxigenic** | 017 | + | + |
|  | **CDL** | CdE-36-CDL | - | **Toxigenic** | 017 | + | + |
| **37** | CAE | CdE-37 | - | **Toxigenic** | 017 | + | + |
|  | **CDL** | CdE-37-CDL | - | **Toxigenic** | 017 | + | + |
| 29* | CAE | CdE-29 | + | Non-Toxigenic | 439 | + (weak) | - |
|  | **CDL** | CdE-29-CDL | - | **Toxigenic** | 005 | + (weak) | - |
| 42 | CAE | CdE-42 | + | **Toxigenic** | 020 | - | - |
|  | **CDL** | CdE-42-CDL | + | **Toxigenic** | 020 | - | - |
| 43 | CAE | CdE-43 | - | **Toxigenic** | 020 | + | - |
|  | **CDL** | CdE-43-CDL | - | **Toxigenic** | 020 | + | - |
| 53 | CAE | CdE-53 | + | **Toxigenic** | 078 | - | - |
|  | **CDL** | none | NA | NA | NA | - | - |
| 54 | CAE | none | NA | NA | NA | - | - |
|  | **CDL** | not preserved | - | **Toxigenic** | ND | - | - |
| 57 | CAE | CdE-57 | - | **Toxigenic** | 020 | - | - |
|  | **CDL** | none | NA | NA | NA | - | - |
| 78 | CAE | CdE-78 | + | **Toxigenic** | 020 | - | - |
|  | **CDL** | none | NA | NA | NA | - | - |
| 81 | CAE | none | NA | NA | NA | - | - |
|  | **CDL** | CdE-81-CDL | + | **Toxigenic** | FR227 | - | - |
| 1 | CAE | CdE-1 | + | **Toxigenic** | AI-53 | - | - |
| 2 | CAE | CdE-2 | + | **Toxigenic** | 005 | + (weak) | - |
| 9 | CAE | CdE-9 | - | **Toxigenic** | 012 | + | - |
| 12 | CAE | CdE-12 | - | **Toxigenic** | 017 | - | - |
| 19 | CAE | CdE-19 | - | **Toxigenic** | 126 | - | - |
| 85 | CAE | CdE-85 | - | **Toxigenic** | 012 | + | - |
| 90 | CAE | CdE-90 | + | **Toxigenic** | 005 | - | - |
| 92 | CAE | CdE-92 | - | **Toxigenic** | 017 | - | + (weak) |
| 18 | CAE | CdE-18 | - | Non-toxigenic | 009 | + | - |
|  | **CDL** | CdE-18-CDL | - | Non-toxigenic | 009 | + | - |
| 48 | CAE | none | NA | NA | NA | - | - |
|  | **CDL** | not preserved | - | Non-toxigenic | ND | - | - |
| 60 | CAE | CdE-60 | - | Non-toxigenic | 009 | + | - |
|  | **CDL** | CdE-60-CDL | - | Non-toxigenic | 009 | + | - |
| 61 | CAE | none | NA | NA | NA | - | - |
|  | **CDL** | CdE-61-CDL | + | Non-toxigenic | 009 | + | - |
| 5 | CAE | CdE-5 | - | Non-toxigenic | 009 | + | - |
| 21 | CAE | CdE-21 | - | Non-toxigenic | 009 | - | - |
| 91 | CAE | CdE-91 | - | Non-toxigenic | 035 | + | - |

Free toxins and GDH were detected in the two samples of four animals (#3, #32, #36 and #37). These animals displayed signs of diarrhea (CDL in the colon). The only identified enteropathogen in their intestinal contents was a toxigenic *C. difficile* (ribotype 126 or 017) and it had expressed its virulence (Table 3, Figure 2). These data were highly suggestive of animal CDI. Sixteen Equidae in which no toxins were detected (Table 3) were carriers of a toxigenic *C. difficile*. Half of them showed no diarrhea signs (#1, #2, #9, #12, #19, #85, #90 and #92), suggesting asymptomatic carriage. For almost all others (except #57), diarrhea signs might be due to another intestinal pathogen (*C. perfringens* in #43, *C. perfringens* or *P. sordellii* in #29 and #42, *C. perfringens*, *P. sordellii* or a rotavirus in #53, and a parasite in #78) or to a non-infectious intestinal disease (torsion of the large colon in #54 and intestinal lymphoma in #81) (Table S1).

**Figure 2.**
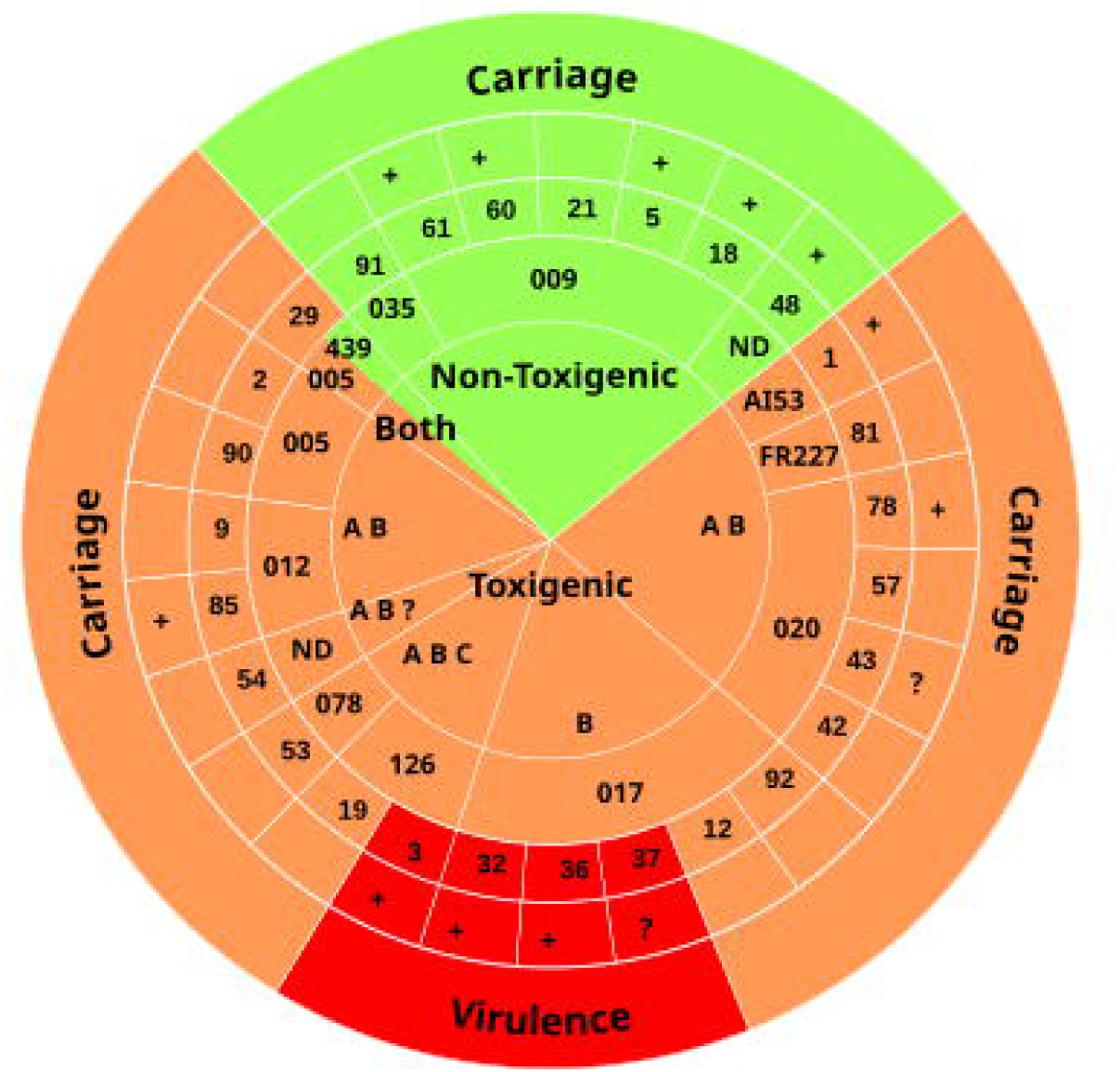
*C. difficile* virulence or carriage in *C. difficile*-positive Equidae. All *C. difficile*-positive animals are shown as a wedge of a pie chart. For each positive animal, the pathogenicity, ribotype and virulence of *C. difficile* are indicated in four concentric rings. First (core): *C. difficile* pathogenicity (Table 2). Non-toxigenic and toxigenic strains are indicated on a green or orange background, respectively. ‘Both’ is indicated when a non-toxigenic strain and a toxigenic strain were co-colonizing the same animal (#29, bi-colored wedge). The toxin gene profile of toxigenic strains is specified using the following abbreviations: A for *tcdA^+^*, B for *tcdB^+^*, C for *cdt^+^* and ‘?’ when the presence of *cdt* genes has not been tested. Second: *C. difficile* ribotype (according to WEBRIBO database) (Table 2). Third: Animal identification number. Fourth: Antibiotic treatment in the two-three weeks before death Fifth (outer ring): Virulence (Toxin production, Table 3) or carriage. Virulence, carriage of a toxigenic strain and carriage of a non-toxigenic strain are on a red, orange and green background, respectively.

In Equidae displaying a toxigenic *C. difficile*, the subpopulations displaying *C. difficile* toxins or not were similar according to sex, age and non-pathognomonic signs of endo-enterotoxaemia or diarrhea, but they significantly differed according to breed and antibiotic treatment (Table 1). Three out of four toxin-positive animals (all Trotters) compared to three out of sixteen carriers of a toxigenic *C. difficile* had received antibiotics (Table S1, Table 1). Antibiotics treatments could therefore have increased the risk for Equidae to develop CDI.

### Cause of animal death

We examined the cause of death of the four toxin-positive animals (Table S1). The fulminant death of animal #3, a 15-days-old foal, was unrelated to *C. difficile* and due to small intestine intussusception (Table S1). Animal #37 died from multiple (respiratory, musculoskeletal and digestive) infections (including endo-enterotoxaemia), probably due to *Klebsiella pneumoniae*, identified in lungs, lung abscess and hock muscle, and/or *C. difficile* (Table S1).

For the last two animals, #32 and #36, death was due to endo-enterotoxaemia (Table S1). Considering the presence of both *C. difficile* and its virulence factors (Table 3), *C. difficile,* was the only identified and most probable cause of death (Table S1). The resulting prevalence of *C. difficile* as the cause of animal death (2%) was of the same range as the previously measured ones (0%-5%). Of note, ribotype 017 was implicated in both cases and probably also in one case in 2018: animal #0, which had died from endo-enterotoxaemia after an antibiotic treatment, and which displayed signs of diarrhea and *C. difficile* as the only enteropathogen (Table 2, Table S1).

## Discussion

### *C. difficile* prevalence and diversity

Here, we evaluated the presence of *C. difficile* in Equidae necropsied in France for the first time. *C. difficile* prevalence in these Equidae having died from diverse, mainly non-intestinal diseases (Table S1) was 27%. This level was similar to, and in some cases lower than, the reported prevalence in European live horses: 4-33% for healthy animals and 5-63% for sick animals with intestinal illnesses (9).

*C. difficile* strains from Equidae belonged to eleven already described ribotypes (9, 32). The historical epidemic ribotype 027 (1), described in horses (19), was not identified here, probably because of the changing *C. difficile* epidemiology and 027 prevalence decrease (32–34). Most ribotypes of CloDifEqui toxigenic strains are common in humans: ribotypes 020, 078, 126 and 005 were in the Top 10 ribotypes in European patients in 2013-2014, and ribotypes 017 and 012 in the Top 20 (32). In Czech horses, toxigenic strains were of five ribotypes (23), including only one in the Top 20 (012) (32). French Equidae could therefore contribute to the dissemination of diverse, clinically important and prevalent ribotypes.

Most ribotypes of CloDifEqui toxigenic strains are also common in animals (9, 11). Ribotypes 126, 020 and 012 had been found in farm animals (9). Ribotype 078, known as prevalent in animals, notably pigs (9), was relatively uncommon here (N=1 isolate), but the phylogenetically close ribotype 126 was represented (N=3) (Table 2) (35). Of note, the ribotypes of most CloDifEqui toxigenic strains have been detected in very diverse sources in Europe, including molluscs (005, 012, 017, 020, 078, 126 and AI-53) (36), potatoes and vegetables (012, 020, 078 and 126) (11, 37). The ubiquity and prevalence of ribotypes identified here suggested that French Equidae could both be contaminated from different sources including humans and contribute to the dissemination of pathogenic *C. difficile* and their transmission to other hosts, notably humans. Equidae could play a role in *C. difficile* circulation and involvement in One Health (19).

### Epidemiology

To our knowledge, this is the first analysis of *C. difficile* ribotypes in animals in France. It was made possible by the registration of detailed animal data and the recovery of animal samples during necropsy, and then by the preservation of ordered collections of animal samples and corresponding *C. difficile* isolates. Following the *ante-mortem* circulation of ribotypes is of great interest from an epidemiological point of view. In two cases, Equidae of the same stud farm were found to share *C. difficile* strains of the same ribotype. This finding indicated that these strains were phylogenetically close, and even raised the possibility that they could form part of a single clonal lineage present in the premises. Each premises could represent a transmission cluster, with a common environmental source contaminating animals or animals contaminating each other. Genome-wide investigations are under progress to fully understand the phylogeny of *C. difficile* strains belonging to each of the ribotypes identified here and their putative transmission between the animals under study.

### Toxin production

After PCR amplification of toxin genes, a method of high sensitivity and negative prediction value, we performed toxin detection, a method of high specificity and positive prediction value, as recommended for CDI diagnosis in European patients (3), but rarely performed in animals including Equidae (38–41). Four animals with both diarrhea signs and a toxigenic *C. difficile* were positive for toxins, suggesting virulence *in vivo* and CDI. *C. difficile* carriers represented 23% of the population (Table 3, Figure 2), a high and previously overlooked level.

In Equidae, *C. difficile* presence (Table 1, Table S1) was significantly correlated to animal signs of diarrhea or endo-enterotoxaemia. In *C. difficile* carriers, in the absence of toxins, these signs were independent from *C. difficile* and nevertheless more frequent than in negative animals (12/23 versus 15/75). Of note, another enteropathogen (bacterium, virus, parasite) was also more frequently identified in *C. difficile* carriers than in negative animals (7/23 versus 9/73) (Table S1), suggesting that it could be responsible for the clinical signs. The most prevalent was *C. perfringens* in four *C. difficile* carriers (Table S1). Host co-colonization or co-infection by *C. difficile* and *C. perfringens* have been described in horses (42, 43), dogs (44), pigs (45) and humans (46).

In this study, the most complex case was animal #29, which displayed three clostridial enteropathogens and two *C. difficile* strains, one toxigenic and the other non-toxigenic (Table S1, Table 2). Animal co-colonization by a toxigenic strain and a non-toxigenic one, notably of ribotype 009, has been reported in companion animals (47). Non-toxigenic strains can prevent CDI or reduce its severity in animal models (48) and a ribotype 009 strain can protect model and commercial piglets (45). Whether CloDifEqui non-toxigenic strains, especially of ribotype 009, could provide protection against a toxigenic strain would require further investigations.

*C. difficile* had produced toxins in four Trotters and at least three had received antibiotics (Table 1, Table S1). Antibiotic treatments could have favored microbiota dysbiosis, *C. difficile* outgrowth and virulence. Of note, among the eight *C. difficile*-positive animals having received antibiotics, five displayed non-toxigenic strains unable to cause disease.

*C. difficile* strains that had produced toxins in Equidae were of ribotypes 126 and 017. Ribotype 126 was the sixth most prevalent ribotype in European patients in 2013-2014 (32). It is also, together with its close relative 078, prevalent in production animals, notably pigs (9, 49), and in biogas plants using animal manure in another French region, Brittany (28). Ribotype 017, which is prevalent in Asia (50), is common in European patients (32) but infrequent and sometimes undetected in French ones (32, 34). As ribotype 017 is also uncommon in animals (9, 50) and was not detected in biogas plants in Brittany (28), its prevalence in French Equidae (5%), and especially those with a toxigenic *C. difficile* (25%) was noticeable. Moreover, its virulence in three close animals (Figure 1) within a short period (Table S1) raised the possibility of a local infection cluster.

In conclusion, our study contributed to a better understanding of the epidemiology of *C. difficile,* by showing the proximity of animals sharing the same ribotype in a small geographical area and sometimes even in a single premises. Of note, we found that 20% of French Equidae necropsied in Normandy are able to disseminate clinically important ribotypes of *C. difficile*. This high, previously unnoticed level of Equidae contamination by toxigenic *C. difficile* would deserve careful monitoring. Our study also revealed that ribotype 017, which is common but relatively infrequent in French hospitals, was prevalent in the studied equine population (5%) and could be virulent (3%) and even cause animal death (2%). These findings raised concerns about the surveillance and control of this ribotype, for the health of Equidae and possibly of the human community. Finally, our study contributed to a better understanding of *C. difficile* from a One Health perspective.

## Statements

### Authorship contribution statement

SP and IP designed the whole project, submitted it to IFCE for funding. JT, NF and MBR performed animal necropsies and intestinal content sampling, registered and analyzed *post-mortem* data. JT established the cause of animal death. KM performed bacteriological and molecular analyses to identify pathogens. SP, KM, FD, SK, LB and IP searched and isolated *C. difficile* strains and performed the primary molecular characterization of strains. FB performed final molecular characterization and free toxin detection. SP and IP analyzed all data. IP wrote the manuscript and SP, JT, FB, FD, SK and IP edited it. All authors read the manuscript and agreed to submit it.

### Data availability

All data are available in the manuscript.

## Acknowledgements

We deeply thank Lydia Baudet (ANSES, Normandy Laboratory for Animal Health, Physiopathology and Epidemiology of Equine Diseases Unit, Goustranville, France), Rabab Syed Zaidi and Lina Ma (CNR *C. difficile*, Hôpital Saint Antoine, Paris, France) for excellent technical assistance. We are grateful to Dr Pierre Nicolas (INRAE, MAIAGE, Jouy-en-Josas, France) for providing helpful insights and reviewing statistical analysis.

