## Supplementary Table 1 for "*Clostridioides difficile* in Equidae necropsied in northwestern France, between 2019 and 2021"

**Supplementary Table 1: Animal Data**

| Animal Id. | Year | Month | Sex | Age range | Breed | French region | Latitude, Longitude | Antibiotic treatment | Hospitalisation before death | Suspision endo-enterotoxæmia | Bacteriological analysis at the veterinary request in case of *post-mortem* observed signs of infection (independent from the systematic *C. difficile* testing) | Cause of death | Samples analysed for this study | *C. difficile* presence (+) and pathogenicity or absence (-) |
|---|---|---|---|---|---|---|---|---|---|---|---|---|---|---|
| **Systematic study 2019-2021** | | | | | | | | | | | | | | |
| **1** | 2019 | May | Male | Foal | Irish Cob | Normandy | 49.1547222,0.3213889 | Yes | No | No | Lung, Liver, kidney: non-hemolytic *E. coli* and *Enterococcus* spp. | Asphyxia by compression of the upper airways (strangles) | Caecum | + toxigenic (tcdA, tcdB) |
| **2** | 2019 | May | Gelding | Adult | Lusitano | Normandy | 49.2426447,-0.8711322 | No | No | No | No bacteriological analysis performed | Anoxia/hypoxia, not pathognomonic. | Caecum | + toxigenic (tcdA, tcdB) |
| **3** | 2019 | May | Male | Foal | French Trotter | Normandy | 49.1458396,0.0005135 | Yes | Yes | No | Caecum, folded colon, small intestine: non-hemolytic *E. coli, Enterococcus faecalis* | Small intestinal intussusception | Caecum, Floating and folded colons | + toxigenic (tcdA, tcdB, cdt) |
| **4** | 2019 | May | Female | Foal | Thoroughbred | Normandy | | No | No | No | Placenta, Lung, Liver, kidney: *Pasteurella pneumotropica,* non-hemolytic *E. coli* , *Enterococcus* spp. | *Pasteurella pneumotropica* infection | Caecum | - |
| **5** | 2019 | May | Female | Foal | Thoroughbred | Normandy | 49.2524182,0.0591629 | Yes | No | No | Lung, lung abscess: *Rhodococcus equi, Enterococcus* spp. | Interstitial pneumonia | Caecum | + non-toxigenic |
| **6** | 2019 | May | Male | Foal | Thoroughbred | Normandy | | Yes | Yes | No | Lung, liver, kidney: *Enterococcus* spp. in addition to non-hemolytic *E. coli* in lung; small intestine content: non-hemolytic *E. coli, Enterococcus faecalis, Staphylococcus xylosus* | Not pathognomonic | Caecum | - |
| **7** | 2019 | May | Female | Adult | French Trotter | Normandy | | No | No | No | No bacteriological analysis performed | Multiparcellar fracture of the cranial | Caecum | - |
| **8** | 2019 | May | Female | Foal | Irish Cob | Normandy | | No | No | No | Placenta, lung, Liver, kidney: *Aeromonas salmonicida, Enterococcus* spp., non-hemolytic *E. coli* in addition to group I *Bacillus* spp., *Staphylococcus xylosus* in placenta; small intestine content: non-hemolytic *E. coli, Enterococcus faecalis, Aeromonas salmonicida, Staphylococcus xylosus* | Systemic infection, degenerative myc | Caecum | - |
| **9** | 2019 | May | Female | Foal | Thoroughbred | Normandy | 48.7069317,0.2196789 | No | No | No | Lung, Liver, kidney: non-hemolytic *E. coli* and *Enterococcus* spp. | Not pathognomonic | Caecum | + toxigenic (tcdA, tcdB) |
| **10** | 2019 | May | Male | Foal | Thoroughbred | Normandy | | Yes | Yes | No | Lung: non-hemolytic *E. coli* | Interstitial pneumonia | Caecum | - |
| **11** | 2019 | June | Female | Adult | Trotter | Normandy | | No | Yes | No | No bacteriological analysis performed | Not pathognomonic | Caecum | - |
| **12** | 2019 | June | Female | Adult | Thoroughbred | Normandy | 48.949829,0.1065658 | No | No | No | No bacteriological analysis performed | Multiparcellar fracture of the left tibi | Caecum | + toxigenic (tcdB) |
| **13** | 2019 | June | Female | Adult | French Trotter | Normandy | | No | No | No | No bacteriological analysis performed | Torsion of the ascending colon & associated circulatory disorders | Caecum, Folded colon | - |
| **14** | 2019 | June | Gelding | Yearlir | Thoroughbred | Normandy | | No | Yes | No | No bacteriological analysis performed | Potential upper cervical trauma | Caecum | - |
| **15** | 2019 | June | Female | Adult | French Trotter | Normandy | | No | No | No | No bacteriological analysis performed | Damaged spinal cord (wobbler) | Caecum | - |
| **16** | 2019 | June | Female | Yearlir | Arabian horse | Normandy | | Yes | No | No | Small intestine content: non-hemolytic *E. coli, Enterococcus faecalis, Proteus mirabilis* ; caecal wall, caecal lymphatic node: *Rhodococcus equi* , non-hemolytic *E. coli, Enterococcus* spp. | *Rhodococcus equi* infection | Caecum, Folded colon | - |
| **17** | 2019 | June | Female | Yearlir | French Trotter | Normandy | | No | No | No | No bacteriological analysis performed | Circulatory & respiratory disorders | Caecum | - |
| **18** | 2019 | June | Female | Foal | Thoroughbred | Normandy | 49.2524182,0.0591629 | Yes | No | Yes | Lung, caudal lobes: non-hemolytic *E. coli, Enterococcus* spp., *Rhodococcus equi* ; pool of digestive contents: non-hemolytic *E. coli, Enterococcus faecalis* | *Rhodococcus equi* infection & endo-enterotoxemia | Caecum, Floating and folded colons | + non-toxigenic |
| **19** | 2019 | June | Female | Young | Thoroughbred | Normandy | 49.3391667,0.0313889 | No | No | Yes | Small intestine content: non-hemolytic *E. coli, Enterococcus faecalis* | Severe hemorrhagic enteritis | Caecum | + toxigenic (tcdA, tcdB, cdt) |
| **20** | 2019 | June | Male | Foal | French Saddlet | Normandy | | No | No | Yes | Lung, Liver, kidney: *Salmonella* spp, *Streptococcus zooepidemicus* , non-hemolytic *E. coli, Enterococcus* spp.; caecum, Folded colon, small intestine: *Salmonella* spp, non-hemolytic *E. coli, Enterococcus faecalis* | Endo-enterotoxemia | Caecum, Floating and folded colons | - |
| **21** | 2019 | June | Female | Foal | Thoroughbred | Normandy | 49.2524182,0.0591629 | No | Yes | No | Lung: *Streptococcus zooepidemicus, Enterococcus* spp.; stifles: *Streptococcus zooepidemicus, Enterococcus* spp., non-hemolytic *E. coli, Staphylococcus xylosus* | Severe stifle disease and pneumonia | Caecum | + non-toxigenic |
| **22** | 2019 | June | Male | Foal | French Saddlet | Normandy | | No | No | No | No bacteriological analysis performed | Small intestinal volvulus | Caecum | - |
| **23** | 2019 | June | Female | Adult | French Trotter | Normandy | | Unknowr | No | No | Liver, liver abscesses: non-hemolytic *E. coli, Streptococcus equi, Enterococcus* spp. | Hepatopathy | Caecum | - |

| | | | | | | | | | | | | | | |
|---|---|---|---|---|---|---|---|---|---|---|---|---|---|---|
| **24** | 2019 | July | Female | Foal | Thoroughbred | Normandy | | Yes | Yes | Yes | Lung: *Streptococcus zooepidemicus,* non-hemolytic *E. coli, Enterococcus* spp.; pool of digestive contents: *Streptococcus zooepidemicus* , non-hemolytic *E. coli, Enterococcus faecalis* | Endo-enterotoxemia & acute bronchopneumonia | Caecum, Floating and folded colons | - |
| **25** | 2019 | July | Female | Adult | Thoroughbred | Normandy | | No | Yes | No | Guttural pockets: *Actinobacillus equili, Streptococcus zooepidemicus, Enterococcus* spp., *Aspergillus fumigatus* | Guttural pouch mycosis | Caecum | - |
| **26** | 2019 | July | Female | Adult | Saddlebred hor | Normandy | | No | Yes | No | No bacteriological analysis performed | Perforation of the floating colon associated with sero-congestive peritonitis | Caecum | - |
| **27** | 2019 | July | Female | Foal | Arabo-Friesian | Centre-Val De Loire | | Yes | No | No | Lung, Liver, kidney: *Enterococcus* spp. | Coagulation deficiency | Caecum | - |
| **28** | 2019 | July | Male | Foal | French Saddleb | Normandy | | Yes | No | No | No bacteriological analysis performed | Myopathy | Caecum | - |
| **29** | 2019 | July | Female | Adult | French Trotter | Normandy | 49.1458396,0.0005135 | No | Yes | Yes | Intestinal content: non-hemolytic *E. coli, C. difficile, Clostridium perfringens, Paeniclostridium sordelli* | Enteritis associated to distension of the digestive reservoirs | Caecum, Intestine | + both non-toxigenic & toxigenic (tcdA, tcdB), co-colonisation |
| **30** | 2019 | July | Male | Foal | Thoroughbred | Normandy | | Yes | Yes | No | Lung: non-hemolytic *E. coli, Enterococcus* spp.; lung abscess: *Rhodococcus equi, Enterococcus* spp. | Bronchiolo-interstitial pneumonia | Caecum | - |
| **31** | 2019 | July | Female | Adult | French Trotter | Normandy | | No | No | No | No bacteriological analysis performed | Rupture of the spleen | Caecum | - |
| **32** | 2019 | July | Female | Foal | French Trotter | Normandy | 48.8866917,0.3153376 | Yes | No | Yes | Caecum: non-hemolytic *E. coli, C. difficile* | Endo-enterotoxemia | Caecum, Folded colon | + toxigenic (tcdB) |
| **33** | 2019 | July | Male | Foal | Thoroughbred | Normandy | | Yes | No | No | Lung: non-hemolytic *E. coli* ; lung abscess: *Rhodococcus equi* | *Rhodococcus equi* infection | Caecum | - |
| **34** | 2019 | July | Male | Foal | Thoroughbred | Normandy | | No | Yes | No | No bacteriological analysis performed | Cervical trauma, total fracture of the axis tooth | Caecum | - |
| **35** | 2019 | July | Female | Adult | French Trotter | Normandy | | No | No | No | No bacteriological analysis performed | Torsion of the large colon | Caecum, Folded colon | - |
| **36** | 2019 | Augus | Female | Foal | French Trotter | Normandy | 48.8312769,0.2794762 | Yes | No | Yes | Pool of digestive contents: non-hemolytic *E. coli, C. difficile* | Endo-enterotoxemia | Caecum, Floating and folded colons | + toxigenic (tcdB) |
| **37** | 2019 | Augus | Female | Adult | French Trotter | Normandy | 49.1184193,-0.0289157 | Unknowr | Yes | Yes | Lung, lung abscess, left hock muscle, hock: *Klebsiella pneumoniae* , non-hemolytic *E. coli* , *Enterococcus* spp.; pool of digestive content: non-hemolytic *E. coli, Enterococcus faecalis* | Several concomitant infectious pathologies with pulmonary, musculoskeletal & digestive localization | Caecum, Floating and folded colons | + toxigenic (tcdB) |
| **38** | 2019 | Augus | Male | Foal | French Trotter | Normandy | | Yes | Yes | No | Lung: *Rhodococcus equi* , non-hemolytic *E. coli* ; trachea, left hock, gastric lymph node abcesses: *Rhodococcus equi, Enterococcus* spp. | *Rhodococcus equi* infection | Caecum | - |
| **39** | 2019 | Augus | Female | Adult | Thoroughbred | Normandy | | No | Yes | No | No bacteriological analysis performed | Traumatic diaphragmatic rupture complicated by displacement of abdominal viscera | Caecum | - |
| **40** | 2019 | Augus | Female | Yearlir | French Trotter | Normandy | | Unknowr | No | No | No bacteriological analysis performed | Severe osteo-articular lesions of both stifles | Caecum | - |
| **41** | 2019 | Septe | Male | Foal | Thoroughbred | Normandy | | No | Yes | No | No bacteriological analysis performed | Displacement with torsion of the various digestive reservoirs & associated circulatory disorders | Caecum | - |
| **42** | 2019 | Septe | Female | Adult | Pony | Normandy | 49.2180615,-0.0988487 | No | No | Yes | Caecum, folded colon, small intestine: non-hemolytic *E. coli, Aerococcus viridans, Enterococcus faecalis, Candida guillermondii* in addition to *Clostridium perfringens* in caecum and *Clostridium perfringens, Paeniclostridium sordelli* in small intestine; Food: non-hemolytic *E. coli, Aerococcus viridans, Candida guillermondii* | Endo-enterotoxemia | Caecum, Folded colon | + toxigenic (tcdA, tcdB) |
| **43** | 2019 | Septe | Female | Adult | Pony | Normandy | 49.2180615,-0.0988487 | Unknowr | No | Yes | Caecum, folded & floating colons, small intestine: non-hemolytic *E. coli, Aerococcus viridans, Enterococcus faecalis* in addition to *Clostridium perfringens* in caecum, folded & floating colon | Endo-enterotoxemia | Caecum, Floating and folded colons | + toxigenic (tcdA, tcdB) |
| **44** | 2019 | Septe | Female | Foal | French Trotter | Normandy | | No | No | No | No bacteriological analysis performed | Parietal tear of the ileum following massive *Paranoplocephala mamillana* infestation | Caecum | - |

| | | | | | | | | | | | | | | |
|---|---|---|---|---|---|---|---|---|---|---|---|---|---|---|
| **45** | 2019 | Septe | Female | Young | Thoroughbred | Normandy | | Yes | No | No | Brain, marrow, lung, liver: *Staphylococcus aureus, Enterococcus* spp. | Systemic infection, myeloencephalitis | Caecum | - |
| **46** | 2019 | Decen | Female | Foal | Thoroughbred | Normandy | | No | No | No | Small intestine content: non-hemolytic *E. coli, Enterococcus faecalis , Streptococcus zooepidemicus* | Small intestinal perforation & subsequent serofibrinous peritonitis | Caecum | - |
| **47** | 2019 | Decen | Male | Young | French Trotter | Normandy | | Unknowr | No | No | No bacteriological analysis performed | Compressive cervical myelopathy | Caecum | - |
| **48** | 2020 | Fevru | Female | Adult | Thoroughbred | Normandy | 49.3212071,-0.7955478 | Yes | No | Yes | Wound, lung, liver, kidney: non-hemolytic *E. coli, Enterococcus* spp in addition to *Streptococcus zooepidemicus* in lung, liver, kidney; small intestine: non-hemolytic *E. coli, Enterococcus faecalis , C. difficile* | Endo-enterotoxemia | Caecum, Small intestine | + non-toxigenic, not preserved |
| **49** | 2020 | Fevru | Male | Young | French Trotter | Normandy | | Yes | No | No | No bacteriological analysis performed | Severe and advanced valvular endocarditis | Caecum | - |
| **50** | 2020 | May | Female | Yearlir | French Trotter | Normandy | | No | No | No | Guttural pockets, lymphatic node, right mandible, soft palate: *Streptococcus equi* ssp. *equi,* non-hemolytic *E. coli, Enterococcus* spp., *Staphylococcus xylosus* | Asphyxia by compression of the upper airways (strangles) | Caecum | - |
| **51** | 2020 | June | Female | Adult | Thoroughbred | Normandy | | Yes | No | No | abdominal abscess: *Enterococcus* spp. non-hemolytic *E. coli , Streptococcus equisimilis, Staphylococcus aureus* (SARM) | Rupture of the stomach following fibrous adhesions | Caecum | - |
| **52** | 2020 | June | Female | Foal | French Trotter | Normandy | | Unknowr | No | No | Lung abscess: *Rhodococcus equi, Streptococcus zooepidemicus, Serratia odorifera, Enterococcus* spp.; pubis: *Streptococcus zooepidemicus, Enterococcus* spp. | Severe and advanced osteomyelitis of the pelvis | Caecum | - |
| **53** | 2020 | June | Female | Foal | Thoroughbred | Normandy | 49.2505283,0.1331312 | No | Yes | Yes | Digestive content: non-hemolytic *E. coli, Enterococcus faecalis, Clostridium perfringens, Paeniclostridium sordellii* ; Floating colon: Rotavirus | Gastric ulcer perforation | Caecum, Floating and folded colons | + toxigenic (tcdA, tcdB, cdt) |
| **54** | 2020 | June | Male | Young | French Trotter | Normandy | 49.1663797,-0.0561634 | No | No | No | No bacteriological analysis performed | Internal bleeding (mesentery) following acute torsion of large colon | Caecum, Floating and folded colons | + toxigenic (tcdA, tcdB, unknown [*cdt* presence not tested]), not preserved |
| **55** | 2020 | June | Female | Foal | French Trotter | Normandy | | Yes | No | No | Floating colon: Rotavirus; digestive content: non-hemolytic *E. coli, Enterococcus faecalis* | Gastric ulcers perforation following s | Caecum, Intest | - |
| **56** | 2020 | June | Female | Yearlir | French Trotter | Normandy | | No | Yes | No | No bacteriological analysis performed | Cranial floor fracture | Caecum | - |
| **57** | 2020 | June | Female | Foal | French Trotter | Normandy | 49.1663797,-0.0561634 | No | No | No | Small inestine : non-hemolytic *E. coli, Enterococcus faecalis, Proteus mirabilis, Streptococcus zooepidemicus* | Myopathy | Caecum, Folded colon | + toxigenic (tcdA, tcdB) |
| **58** | 2020 | July | Male | Foal | Thoroughbred | Normandy | | Yes | No | No | Lung, lung abscess: *Rhodococcus equi, Enterococcus* spp. | *Rhodococcus equi* infection | Caecum | - |
| **59** | 2020 | July | Female | Foal | Thoroughbred | Normandy | | No | No | No | Floating colon: Rotavirus | Small intestinal volvulus | Caecum, Small intestine | - |
| **60** | 2020 | July | Female | Foal | French Trotter | Normandy | 48.999875,0.0952281 | Yes | Yes | Yes | Intestinal content: Rotavirus | Acute enteritis | Caecum, Folded colon and small intestine | + non-toxigenic (initial detection: toxigenic $tcdB^{+}$) |
| **61** | 2020 | July | Female | Foal | Thoroughbred | Normandy | 49.2524182,0.0591629 | Yes | Yes | No | No bacteriological analysis performed | Small intestinal volvulus & infarction | Caecum, Small intestine | + non-toxigenic (initial detection: toxigenic $tcdA^{+}$, $tcdB^{+}$) |
| **62** | 2020 | July | Male | Foal | Thoroughbred | Normandy | | Unknowr | Yes | No | Lung: non-hemolytic *E. coli, Enterococcus* spp. | Interstitial bronchopneumonia | Caecum | - |
| **63** | 2020 | July | Male | Foal | French Trotter | Normandy | | Yes | No | No | Lung abscess, lymphatic node: *Rhodococcus equi* , non-hemolytic *E. coli, Proteus vulgaris, Enterococcus* spp.; lung: *Rhodococcus equi* , non-hemolytic *E. coli* | *Rhodococcus equi* infection | Caecum, Floating and folded colons | - |
| **64** | 2020 | July | Female | Foal | Thoroughbred | Normandy | | Yes | No | No | Lung: *Rhodococcus equi* ; Lymphatic node: *Rhodococcus equi* , non-hemolytic *E. coli, Enterococcus* spp.; lung abscess: *Rhodococcus equi, Enterococcus s* pp. ; PG: *Rhodococcus equi, Pseudomas aeruginosa, Enterococcus* spp., non-hemolytic *E. coli* | *Rhodococcus equi* infection | Caecum | - |
| **65** | 2020 | July | Male | Yearlir | French Trotter | Normandy | | No | Yes | No | No bacteriological analysis performed | Compressive cervical myelopathy | Caecum | - |
| **66** | 2020 | Augus | Unknow | Foal | Unknown | Unknown | | No | No | Unknown | Unknown | Unknown | Intestine | - |
| **67** | 2020 | Septe | Gelding | Adult | Saddlebred hor | Normandy | | No | No | No | No bacteriological analysis performed | Not pathognomonic | Caecum | - |
| **68** | 2020 | Septe | Gelding | Adult | Pony | Normandy | | No | No | Yes | No bacteriological analysis performed | Endo-enterotoxemia | Caecum | - |

| | | | | | | | | | | | | | | |
|---|---|---|---|---|---|---|---|---|---|---|---|---|---|---|
| **69** | 2020 | Septe | Gelding | Young | Pony | Normandy | | No | No | No | Lung, Liver, kidney: *Pasteurella pneumotropica,* non-hemolytic *E. coli, Enterococcus* spp. in addition to *Streptococcus zooepidemicus* in lung | Systemic infection, interstitial pneumonia | Caecum | - |
| **70** | 2020 | Septe | Gelding | Adult | French Saddleb | Normandy | | Yes | Unknow | No | Endocardium, lung, liver, kidney: non-hemolytic *E. coli, Enterococcus* spp. | Hemorrhagic syndrome | Caecum | - |
| **71** | 2020 | Septe | Female | Adult | French Saddleb | Normandy | | No | Yes | No | No bacteriological analysis performed | Small intestinal volvulus | Caecum | - |
| **72** | 2020 | Octob | Gelding | Foal | Irish Cob | Normandy | | No | No | No | Lung, lymphatic node: *Streptococcus zooepidemicus, Enterococcus* spp. | Compressive cervical myelopathy | Caecum | - |
| **73** | 2020 | Octob | Male | Young | French Trotter | Normandy | | Unknowr | No | No | No bacteriological analysis performed | Atypical myopathy | Caecum | - |
| **74** | 2020 | Nover | Male | Yearlir | Thoroughbred | Normandy | | Yes | No | Yes | Lung, Liver, kidney: non-hemolytic *E. coli, Enterococcus* spp. ; digestive content: *Enterococcus faecalis* , non-hemolytic *E. coli* ; colons: *Clostridium perfringens* | Endo-enterotoxemia | Caecum, Folded colon | - |
| **75** | 2020 | Nover | Male | Foal | French Trotter | Normandy | | No | Yes | No | No bacteriological analysis performed | Degenerative myopathy | Caecum | - |
| **76** | 2020 | Nover | Female | Foal | Thoroughbred | Normandy | | Yes | No | Yes | Lung abscess: *Staphylococcus aureus, Pseudomonas* spp. | Endo-enterotoxemia | Caecum, Folded colon | - |
| **77** | 2020 | Nover | Female | Yearlir | Thoroughbred | Normandy | | No | Yes | No | No bacteriological analysis performed | Pelvic fracture | Caecum | - |
| **78** | 2020 | Nover | Female | Adult | Saddlebred hoı | Normandy | 49.0385627746582,0.7007 | Yes | No | Yes | Absence of *Salmonella* spp. | Larval cyathostomy | Caecum, Folded colon | + toxigenic (tcdA, tcdB) |
| **79** | 2020 | Nover | Female | Adult | Thoroughbred | Normandy | | No | No | No | lymphatic node: *Enterococcus* spp., *Klebsiella pneumoniae* ssp. *ozaenae* | Rupture of the small intestine | Caecum | - |
| **80** | 2020 | Decen | Female | Adult | Donkey | Normandy | | No | No | Yes | Caecocolic content: non-hemolytic *E. coli, Salmonella Enteritidis, Clostridium perfringens* ; folded colon: non-hemolytic *E. coli, Enterococcus faecalis, Streptococcus zooepidemicus* | Endo-enterotoxemia | Caecum, Floating and folded colons | - |
| **81** | 2020 | Decen | Gelding | Adult | Irish Cob | Normandy | 49.1547222,0.3213889 | No | No | Yes | Digestive content: *E.coli* ; small intestine abscess: non-hemolytic *E. coli, Klebsiella pneumoniae, Streptococcus zooepidemicus, Enterococcus faecalis* ; liver: *Enterococcus* spp., *Streptococcus zooepidemicus, Klebsiella pneumoniae* | Intestinal lymphoma | Caecum, Floating and folded colons | + toxigenic (tcdA, tcdB) |
| **82** | 2020 | Decen | Female | Adult | Pony | Normandy | | No | No | No | Rate: *Babesia caballi* | Rupture of the spleen | Caecum | - |
| **83** | 2020 | Decen | Female | Foal | French Saddleb | Normandy | | Yes | No | No | Fetlock: *Aerococcus viridans,* non-hemolytic *E. coli, Enterococcus* spp., *Saccharomyces cerevisiae* | Post-infectious necrotizing systemic vasculitis (hemorrhagic purpura) | Caecum | - |
| **84** | 2021 | Janua | Female | Adult | Thoroughbred | Normandy | | No | No | No | No bacteriological analysis performed | Rupture of the vaginal artery | Caecum | - |
| **85** | 2021 | Janua | Male | Foal | French Saddleb | Normandy | 49.1656553,-1.0074577 | Yes | Yes | No | Paratesticular abscess: non-hemolytic *E. coli, Streptococcus zooepidemicus* | Duodenum ulcer perforation | Caecum | + toxigenic (tcdA, tcdB) |
| **86** | 2021 | Janua | Female | Adult | French Trotter | Brittany | | No | No | No | No bacteriological analysis performed | Grass sickness | Caecum | - |
| **87** | 2021 | Fevru | Female | Adult | Thoroughbred | Normandy | | No | Yes | No | No bacteriological analysis performed | Eosinophilic enterocolitis | Caecum, Folded colon | - |
| **88** | 2021 | Fevru | Female | Adult | Thoroughbred | Normandy | | No | No | No | No bacteriological analysis performed | Dystocic foaling | Caecum | - |
| **89** | 2021 | Fevru | Female | Adult | Thoroughbred | Normandy | | No | Yes | No | No bacteriological analysis performed | Rupture of the uterine artery | Caecum | - |
| **90** | 2021 | March | Female | Yearlir | Thoroughbred | Normandy | 49.0919444,0.1063889 | No | Yes | No | No bacteriological analysis performed | Parietal tear of the ileum following massive *Paranoplocephala mamillana* infestation | Caecum | + toxigenic (tcdA, tcdB) |
| **91** | 2021 | March | Female | Foal | Thoroughbred | Normandy | 48.74993,0.2964154 | No | Yes | No | Lung, Liver, kidney, guttural pockets: *Enterococcus* spp.; small intestine content: non-hemolytic *E. coli, Enterococcus faecalis* | Systemic infection | Caecum | + non-toxigenic |
| **92** | 2021 | April | Female | Foal | Thoroughbred | Normandy | 49.1397709,0.2094533 | No | No | No | Lung, Liver, kidney, right stifle: *Staphylococcus aureus, Streptococcus zooepidemicus* in addition to *Enterococcus* spp. in right stifle | Severe and extensive pleuropneumonia | Caecum | + toxigenic (tcdB) |
| **93** | 2021 | April | Female | Foal | Unknown | Normandy | | No | No | No | *Streptococcus equi* ssp. *equi* | Atypical form of strangles with sero-f | Caecum | - |
| **94** | 2021 | May | Female | Adult | French Trotter | Normandy | | Yes | No | No | *Clostridium perfringens, Paeniclostridium sordellii* | Myonecrosis | Caecum | - |
| **95** | 2021 | June | Female | Foal | Thoroughbred | Normandy | | Yes | Unknow | No | Non-hemolytic *E. coli* | Perforation of the floating colon | Caecum | - |
| **96** | 2021 | June | Female | Adult | Saddlebred hoı | Normandy | | No | No | No | No bacteriological analysis performed | Rhinopneumonia (nervous form) | Caecum | - |
| **97** | 2021 | June | Female | Yearlir | Thoroughbred | Normandy | | Yes | No | No | Small inestine : non-hemolytic *E. coli, Enterococcus faecalis* | Ileocaecal intussusception | Caecum | - |
| **98** | 2021 | June | Female | Foal | Thoroughbred | Normandy | | No | No | No | No bacteriological analysis performed | Rupture of the small intestine due to *Parascaris* | Caecum | - |

| | | | | | | | | | | | | | | |
|---|---|---|---|---|---|---|---|---|---|---|---|---|---|---|
| **99** | 2021 | July | Male | Foal | French chaser | Normandy | | Yes | Yes | No | Lung abscess, caecum, forlded colon: *Rhodococcus equi,* non-hemolytic *E. coli* ; respiratory swab: *Rhodococcus equi, Citrobacter freundii* | *Rhodococcus equi* infection | Caecum | - |
| **100** | 2021 | July | Female | Foal | Thoroughbred | Normandy | | No | Yes | Yes | *Rhodococcus equi* ; pool of digestive content: non-hemolytic *E. coli* ; small interstine: *Clostridium perfringens* | Acute typhlocolitis & *Rhodococcus equi* infection | Caecum, Folded colon | - |
| **Pilot study 2018** | | | | | | | | | | | | | | |
| **0** | 2018 | Septe | Female | Foal | French Saddlebred Horse | Normandy | | Yes | | Yes | Folded colon: *Clostridium difficile,* non-hemolytic *Escherichia coli, Enterococcus faecalis* ; lung, guttural pocket: *Klebsiella pneumoniae* | Endo-enterotoxemia | Folded colon | + toxigenic (tcdB) |

The registered data of necropsied Equidae are shown.
