## Supplementary Table 2 for "*Clostridioides difficile* in Equidae necropsied in northwestern France, between 2019 and 2021"

**Supplementary Table S2: Sequence Type (MLST) of CloDifEqui strains**

| ***C. difficile* strain** | Equidae | | **Allele of housekeeping gene** | | | | | | | **MLST** | Ribotype | Toxin gene profile* | Clade |
|---|---|---|---|---|---|---|---|---|---|---|---|---|---|
| | n° | Digestive content | ***adk*** | ***atpA*** | ***dxr*** | ***glyA*** | ***recA*** | ***sodA*** | ***tpi*** | | | | |
| *Cd* E3 | 3 | CAE | 5 | 8 | 5 | 11 | 9 | 11 | 8 | **11** | 126 | $tcdA^{+}$ $tcdB^{+}$ $cdt^{+}$ | 5 |
| *Cd* E3-CDL | | CDL | 5 | 8 | 5 | 11 | 9 | 11 | 8 | | | | |
| *Cd* E19 | 19 | CAE | 5 | 8 | 5 | 11 | 9 | 11 | 8 | | | | |
| *Cd* E9 | 9 | CAE | 1 | 4 | 7 | 1 | 1 | 3 | 3 | **54** | 012 | $tcdA^{+}$ $tcdB^{+}$ | 1 |
| *Cd* E43 | 43 | CAE | 1 | 1 | 2 | 5 | 1 | 3 | 1 | **2** | 020 | $tcdA^{+}$ $tcdB^{+}$ | 1 |
| *Cd* E43-CDL | | CDL | 1 | 1 | 2 | 5 | 1 | 3 | 1 | | | | |
| *Cd* E12 | 12 | CAE | 3 | 7 | 3 | 8 | 6 | 9 | 11 | **37** | 017 | $tcdB^{+}$ | 4 |
| *Cd* E32 | 32 | CAE | 3 | 7 | 3 | 8 | 6 | 9 | 11 | | | | |
| *Cd* E32-CDL | | CDL | 3 | 7 | 3 | 8 | 6 | 9 | 11 | | | | |
| *Cd* E36 | 36 | CAE | 3 | 7 | 3 | 8 | 6 | 9 | 11 | | | | |
| *Cd* E36-CDL | | CDL | 3 | 7 | 3 | 8 | 6 | 9 | 11 | | | | |
| *Cd* E37-CDL | 37 | CDL | 3 | 7 | 3 | 8 | 6 | 9 | 11 | | | | |
| *Cd* E5 | 5 | CAE | 1 | 1 | 2 | 1 | 1 | 1 | 1 | **3** | 009 | Non Toxigenic | 1 |
| *Cd* E18 | 18 | CAE | 1 | 1 | 2 | 1 | 1 | 1 | 1 | | | | |
| *Cd* E18-CDL | | CDL | 1 | 1 | 2 | 1 | 1 | 1 | 1 | | | | |

* according to multiplex PCR results

The Sequence Type (MLST) of the first isolated strains was established.
