## Supplementary Table 3 for "*Clostridioides difficile* in Equidae necropsied in northwestern France, between 2019 and 2021"

**Supplementary Table S3: Toxin detection in *C. difficile* -negative samples**

| Equidae n° | Digestive content | Number of tested versus recovered contents | *C. difficile* detection | Enzymatic Immuno Assay GDH | Free Toxins (Toxin A Toxin B) |
|---|---|---|---|---|---|
| 20 | CAE | 2/2 | - | - | - |
| | **CDL** | | - | - | - |
| 24 | CAE | 2/2 | - | - | - |
| | **CDL** | | - | - | - |
| 55 | CAE | 2/2 | - | - | - |
| | **CDL** | | - | - | - |
| 59 | CAE | 2/2 | - | - | - |
| | **CDL** | | - | - | - |
| 63 | **CDL** | 2/2 | - | - | - |
| 74 | CAE | 2/2 | - | - | - |
| | **CDL** | | - | - | - |
| 76 | CAE | 2/2 | - | - | - |
| | **CDL** | | - | - | - |
| 80 | CAE | 2/2 | - | - | - |
| | **CDL** | | - | - | - |
| 13 | **CDL** | 1/2 | - | - | - |
| 16 | **CDL** | 1/2 | - | - | - |
| 35 | **CDL** | 1/2 | - | - | - |
| 87 | **CDL** | 1/2 | - | - | - |
| 68 | CAE | 1/1 | - | - | - |

The digestive contents of *C. difficile* -negative animals that have been tested for the presence of toxins as controls are listed.

All digestive contents of *C. difficile* -positive animals, whatever toxin detection result, are presented in Table 3.
